## Supplementary material for "The GlycoPaSER prototype as a real-time N-glycopeptide identification tool based on the PaSER parallel computing platform": Supplemetary Figures

**Figure S1: Common glycopeptide fragments (CID) nomenclature [25]. (A)** The nomenclature for each peptide bond or glycosidic bond fragmentation. Branching order (α,β, ‘, ‘’) are determined by branch mass. Peptide bond cleavage result in b and y fragments while cleavage of the glycosidic bond yields B and Y fragments (capitals) **(B)** Examples of glycopeptide fragments. B ions and their fragments are named oxonium ions, after their oxygen charge carrier.

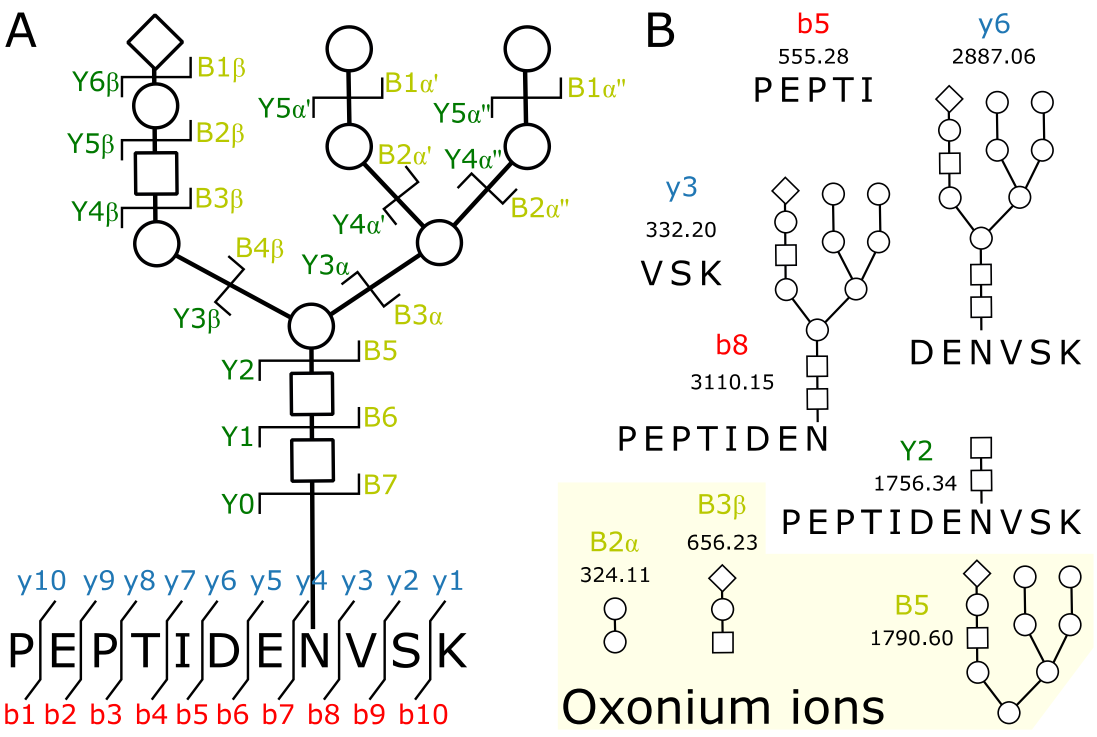

**Figure S2**: **Examples of glycopeptide fragmentation spectra** before (top) and after (bottom) the modification in the glycopeptide decomposer. The oxonium ions and N-glycan core fragmentation pattern matches are highlighted and annotated. p stands for the peptide-moiety mass.

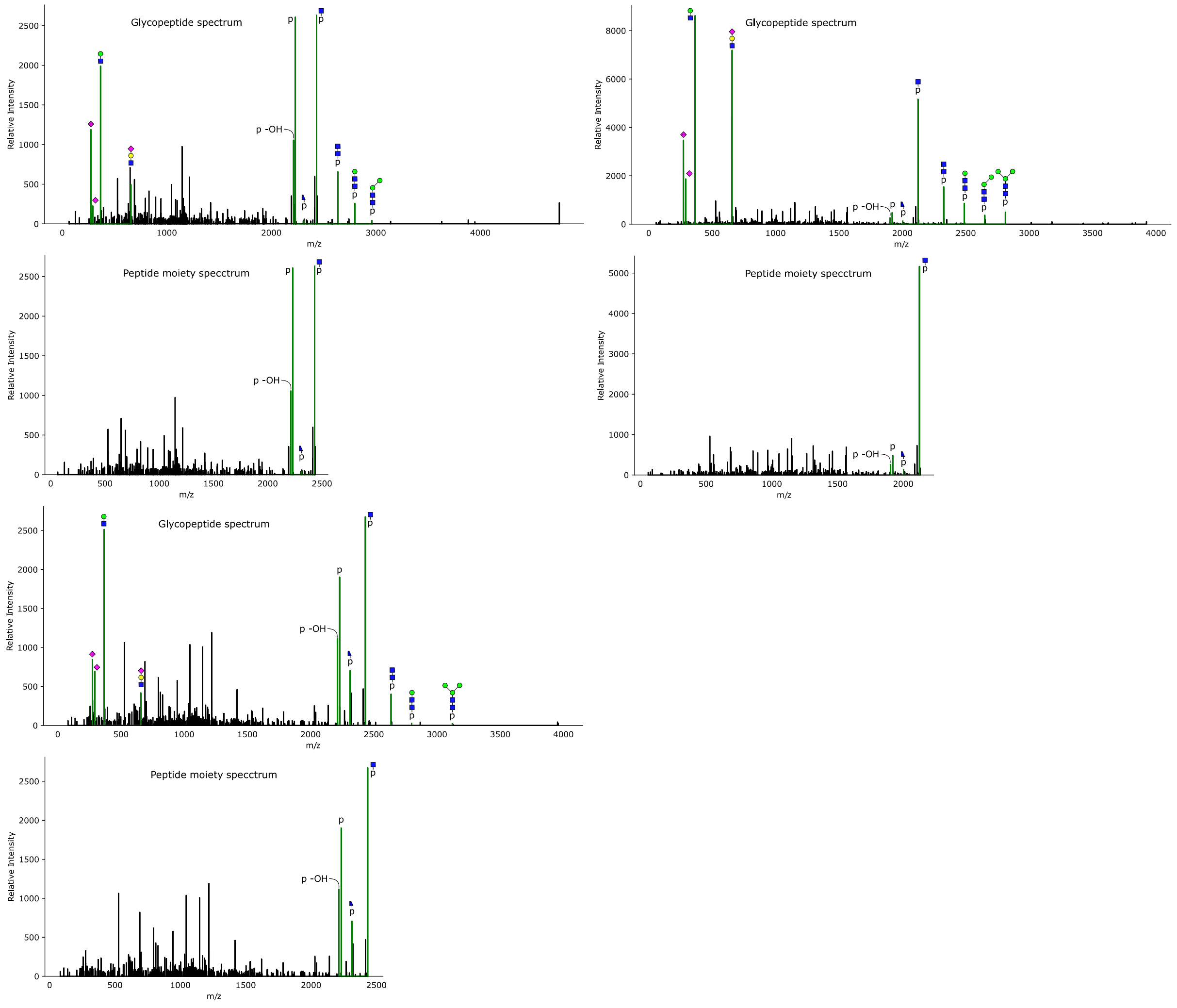

**Figure S3**: **Oxonium ions filter parameter selection. (A)** The relative spectra count for each of the tested oxonium ions. **(B)** The number of spectra with at least X oxonium ions. **(C)** ROC curve for relative intensity sum of oxonium ions (as in Figure 2B), positives are glycopeptides, negatives are non-glycopeptides. The selected threshold is indicated.

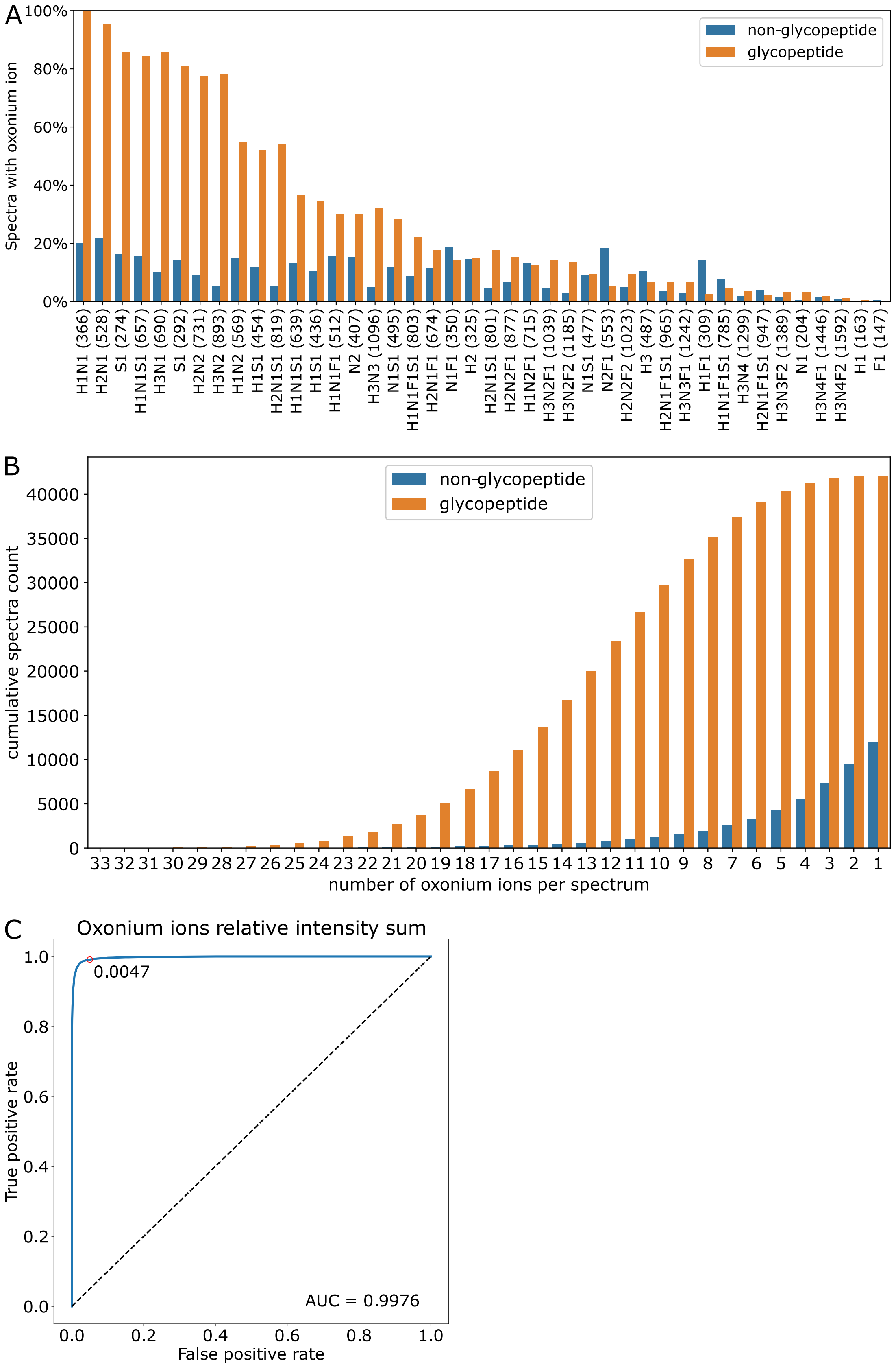

**Figure S4**: Processing time per spectrum for the new GlycoPaSER modules. The average time for glycopeptide decomposer total is shorter than the average time for the pattern finder since the module does not search for the glycopeptide fragmentation pattern in every spectrum.

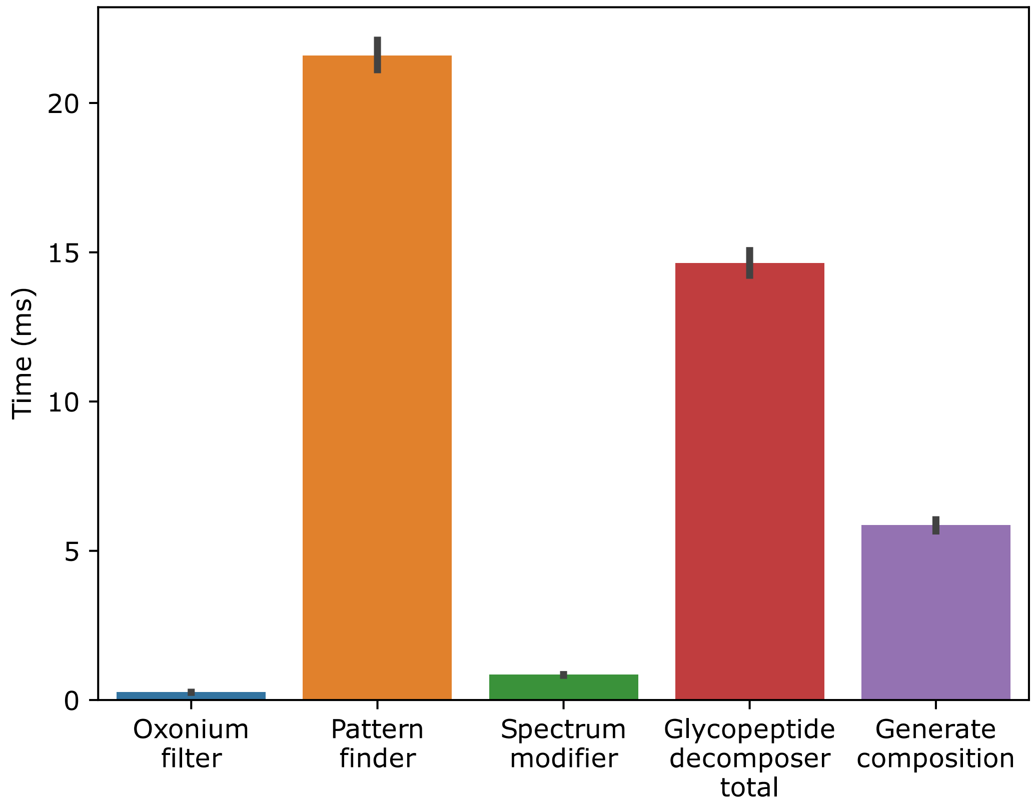

**Figure S5**: Distribution of possible compositions per glycopeptide identification in PaSER.

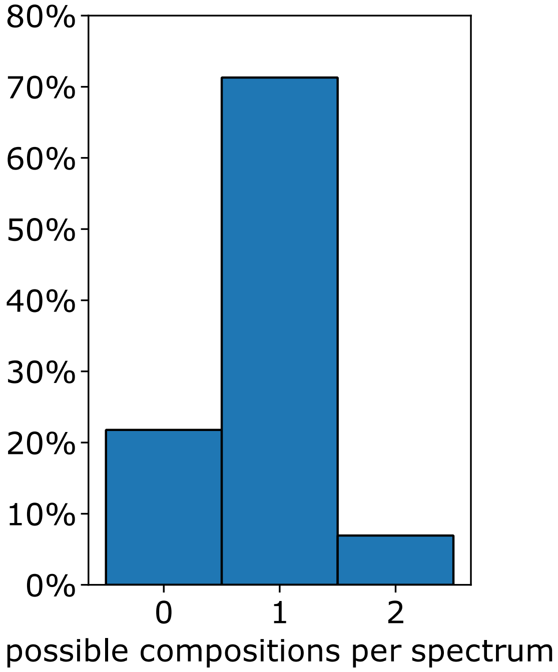


**Figure S6**: Score increase distribution for a file with mixed spectra each from the optimal CE settings compared to the original file acquired with the optimized default CE settings.

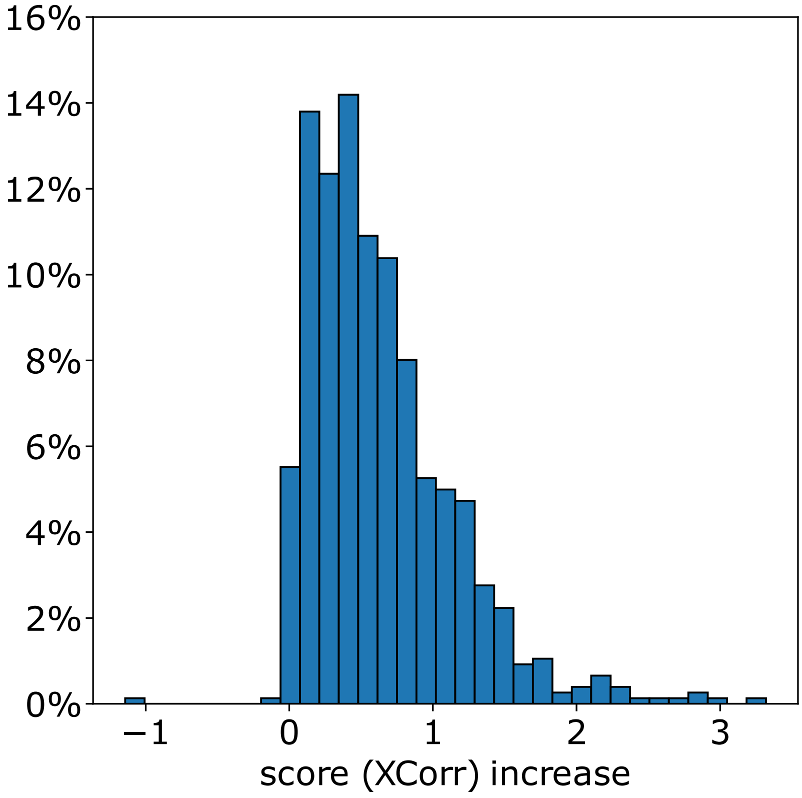
